# Cellpose 2.0: how to train your own model

**DOI:** 10.1101/2022.04.01.486764

**Authors:** Carsen Stringer, Marius Pachitariu

## Abstract

Generalist models for cellular segmentation, like Cellpose, provide good out-of-the-box results for many types of images. However, such models do not allow users to adapt the segmentation style to their specific needs and may perform sub-optimally for test images that are very different from the training images. Here we introduce Cellpose 2.0, a new package which includes an ensemble of diverse pretrained models as well as a human-in-the-loop pipeline for quickly prototyping new specialist models. We show that specialist models pretrained on the Cellpose dataset can achieve state-of-the-art segmentation on new image categories with very little user-provided training data. Models trained on 500-1000 segmented regions-of-interest (ROIs) performed nearly as well as models trained on entire datasets with up to 200,000 ROIs. A human-in-the-loop approach further reduced the required user annotations to 100-200 ROIs, while maintaining state-of-the-art segmentation performance. This approach enables a new generation of specialist segmentation models that can be trained on new image types with only 1-2 hours of user effort. We provide software tools including an annotation GUI, a model zoo and a human-in-the-loop pipeline to facilitate the adoption of Cellpose 2.0.

## Introduction

Biological images of cells are highly diverse due to the combinatorial options provided by various microscopy techniques, tissue types, cell lines, fluorescence labelling etc [1–4]. The available options for image acquisition continue to diversify as advances in biology and microscopy allow for monitoring a larger diversity of cells and signals. This diversity of methods poses a grand challenge to automated segmentation approaches, which have traditionally been developed for specific applications, and fail when applied to novel types of data.

High-performance segmentation methods now exist for several applications [5–9]. These algorithms typically rely on large training datasets of human-labelled images and neural network-based models trained to reproduce these annotations. Such models draw heavy inspiration from the machine vision literature of the last ten years, which is dominated by neural networks. However, neural networks struggle to generalize to out-of-distribution data, i.e. new images that look fundamentally different from anything seen during training. To mitigate this problem, machine vision researchers assemble diverse training datasets, for example by scraping images from the internet or adding perturbations [10, 11]. Computational biologists have tried to replicate this approach by constructing training datasets that were either diverse (Cellpose) or large (TissueNet, LiveCell). Yet even models trained on these datasets can fail on new categories of images (e.g. the Cellpose model on Tissuenet or LiveCell data, Figure 3ac).

Thus, a challenge arises: how can we ensure accurate and adaptable segmentation methods for new types of biological images? Recent studies have suggested novel architectures, new training protocols and image simulation methods for attaining high-performance segmentation with limited training data [12–15]. An alternative approach is provided by interactive machine learning methods. For example, methods like Ilastik allow users to both annotate their data and train models on their own annotations [16]. Another class of interactive approaches known as “human-in-the-loop” start with a small amount of user-segmented data to train an initial, imperfect model. The imperfect model is applied to other images, and the results are corrected by the user. This is the strategy used to annotate the TissueNet dataset, which in total took two human years of crowdsourced work for 14 image categories ([6, 17]). The annotation/retraining process can also be repeated in a loop until the entire dataset has been segmented. This approach has been demonstrated for simple ROIs like nuclei and round cells, which allow for weak annotations like clicks and squiggles ([18, 19]), but not for cells with complex morphologies which require full cytoplasmic segmentation. For example, using an iterative approach, [19] segmented a 3D dataset of nuclei in approximately one month. It is not clear if the human-in-the-loop approach can be accelerated further, and whether it can in fact achieve state-of-the-art performance on cellular images, i.e. human levels of accuracy.

Here we developed algorithmic and software tools for adapting neural network segmentation models to new image categories with very little new training data. We demonstrate that this approach is: a) necessary, because annotation styles can vary dramatically between different annotators; b) efficient, because it only requires a user to segment 500-1000 ROIs offline or 100-200 ROIs with a human-in-the-loop approach; and c) effective, because models created this way have similar accuracy to human experts. We performed these analyses on two large-scale datasets released recently [6, 7] and we used Cellpose, a generalist model for cellular segmentation [5]. We took advantage of these new datasets to develop a model zoo of pretrained models, which can be used as starting points for the human-in-the-loop approach. We also developed a user-friendly pipeline for human-in-the-loop annotation and model retraining. An annotator using our GUI was able to generate state-of-the-art models in 1-2 hours per category.

## Results

### Human annotators use diverse segmentation styles

The original Cellpose is a generalist model that can segment a wide variety of cellular images [5]. We gradually added more data to this model based on user contributions, and we wanted to also add data from the TissueNet and LiveCell datasets [6, 7]. However, we noticed that many of the annotation styles in the new datasets were conflicting with the original Cellpose segmentation style. For example, nuclei were not segmented in the Cellpose dataset if they were missing a cytoplasm or membrane label (Figure 1ai), but they were always labelled in the TissueNet dataset (Figure 1bi, Figure 1biii). Processes that were diffuse were not segmented in the Cellpose dataset (Figure 1aii) but they were always segmented in the LiveCell dataset (Figure 1civ). The outlines in the Cellpose dataset were drawn to include the entire cytoplasm of each cell, often biased towards the exterior of the cell (Figure 1aiii). Some TissueNet categories also included the entire cytoplasm (Figure 1bi), but others excluded portions of the cytoplasm (Figure 1bii, Figure 1biii), or even focused exclusively on the nucleus (Figure 1biv). Finally, areas of high density and low confidence were nonetheless given annotations in the Cellpose dataset and in some LiveCell categories (Figure 1aiv, Figure 1ci), while they were often not segmented in other LiveCell categories (Figure 1cii, Figure 1ciii, Figure 1civ).

**Figure 1:**
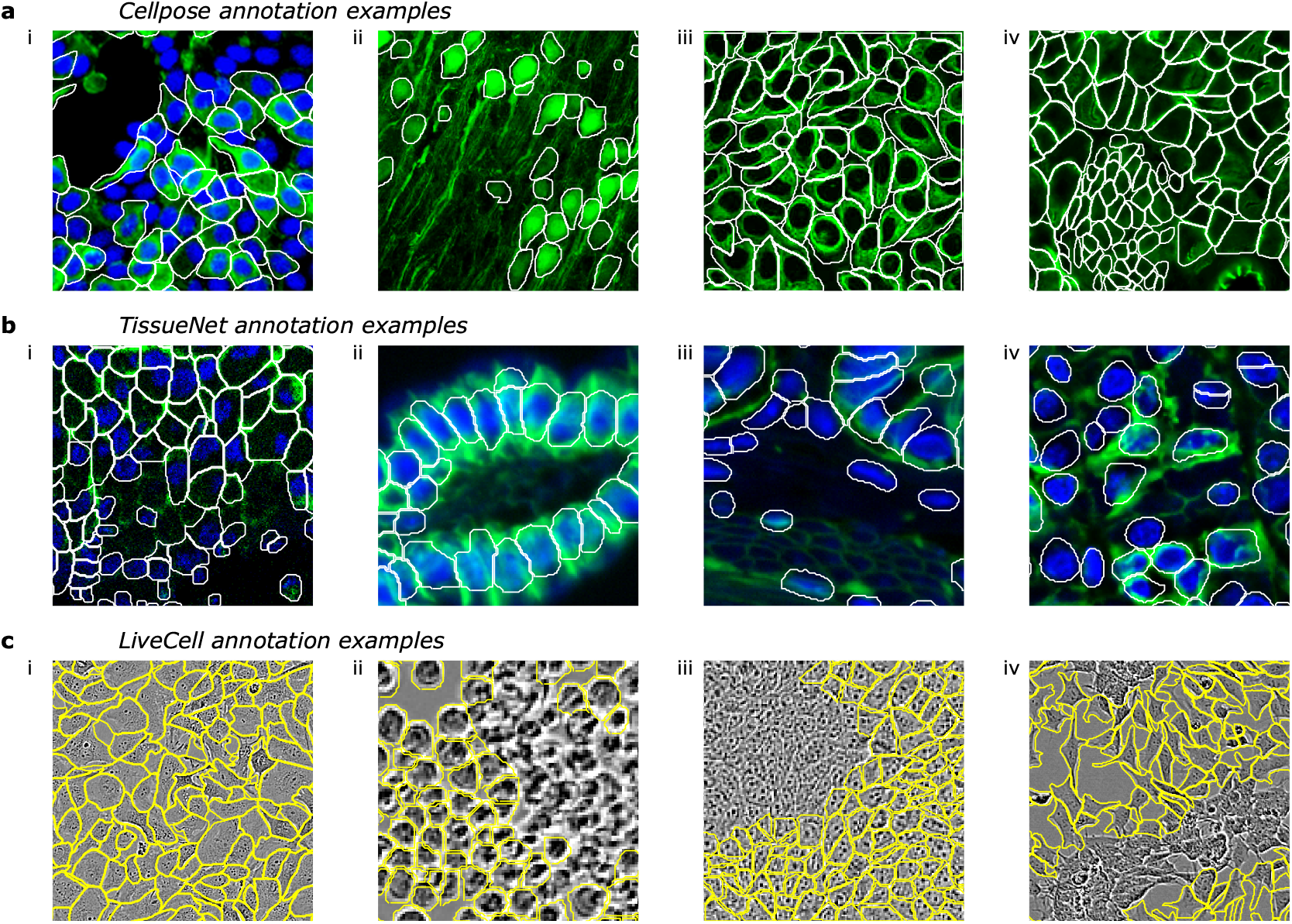
Diverse annotation styles across ground truth datasets. These are examples of images which the human annotators chose to segment a certain way, where another equally valid segmentation style exists. All these examples were chosen to be representative of large categories of images in their respective datasets. **a**, Annotation examples from the Cellpose dataset. From left to right, these show: i) nuclei without cytoplasm are not labelled, ii) diffuse processes are not labelled, iii) outlines biased towards outside of cells, iv) dense areas with unclear boundaries are nonetheless segmented. **b**, Annotation examples from the TissueNet dataset. These illustrate: i) outlines follow membrane/cytoplasm for some image types, and include nuclei without a green channel label, ii) outlines do not follow cytoplasm for other image types, iii) slightly out of focus cells are not segmented, iv) outlines drawn just around nucleus for some image types. **c**, Annotation examples from the LiveCell dataset. These illustrate: i) dense labelling for some image types, ii) no labelling in dense areas for other image types, iii) same as ii, iv) no labelling in some image areas for unknown reasons.

### Creating a model zoo for Cellpose

These examples of conflicting segmentations were representative of large classes of images from across all three datasets. Given this variation in segmentation styles, we reasoned that a single global model may not perform best on all images. Thus, we decided to create an ensemble of models which a user can select between and evaluate on their own data. This would be similar to the concept of a “model zoo” available for other machine learning tasks [20–22].

To synthesize a small ensemble of models, we developed a clustering procedure which groups images together based on their segmentation style (see also [12]). As a marker of the segmentation style we used the style vectors from the Cellpose model [5, 23]. This representation summarizes the style of an image with a “style vector” computed at the most downsampled level of the neural network. The style vector is then broadcast broadly to all further computations, directly affecting the segmentation style of the network. We took the style vectors for all images and clustered them into nine different classes using the Leiden algorithm, illustrated on a t-SNE plot in Figure 2a [24, 25]. For each class, we assigned it a name based on the most common image type included in that class. There were four image classes composed mainly of fluorescent cell images (CP, TN1, TN2, TN3), four classes composed mainly of phase contrast images (LC1, LC2, LC3, LC4) and a ninth class including a wide variety of images (CPx) (Figure 2b). For each cluster, we trained a separate Cellpose model. At test time, new images were co-clustered with the predetermined segmentation styles and assigned to one of the nine clusters. Then the specific model trained on that class was used to segment the image. The ensemble of models significantly outperformed a single global model (Figure 2c). All image classes had improvements in the range of 0.01-0.06 for the AP score, with the largest improvements observed at higher intersection-over-union (IoU) thresholds, and for the most diverse image class (CPx).

**Figure 2:**
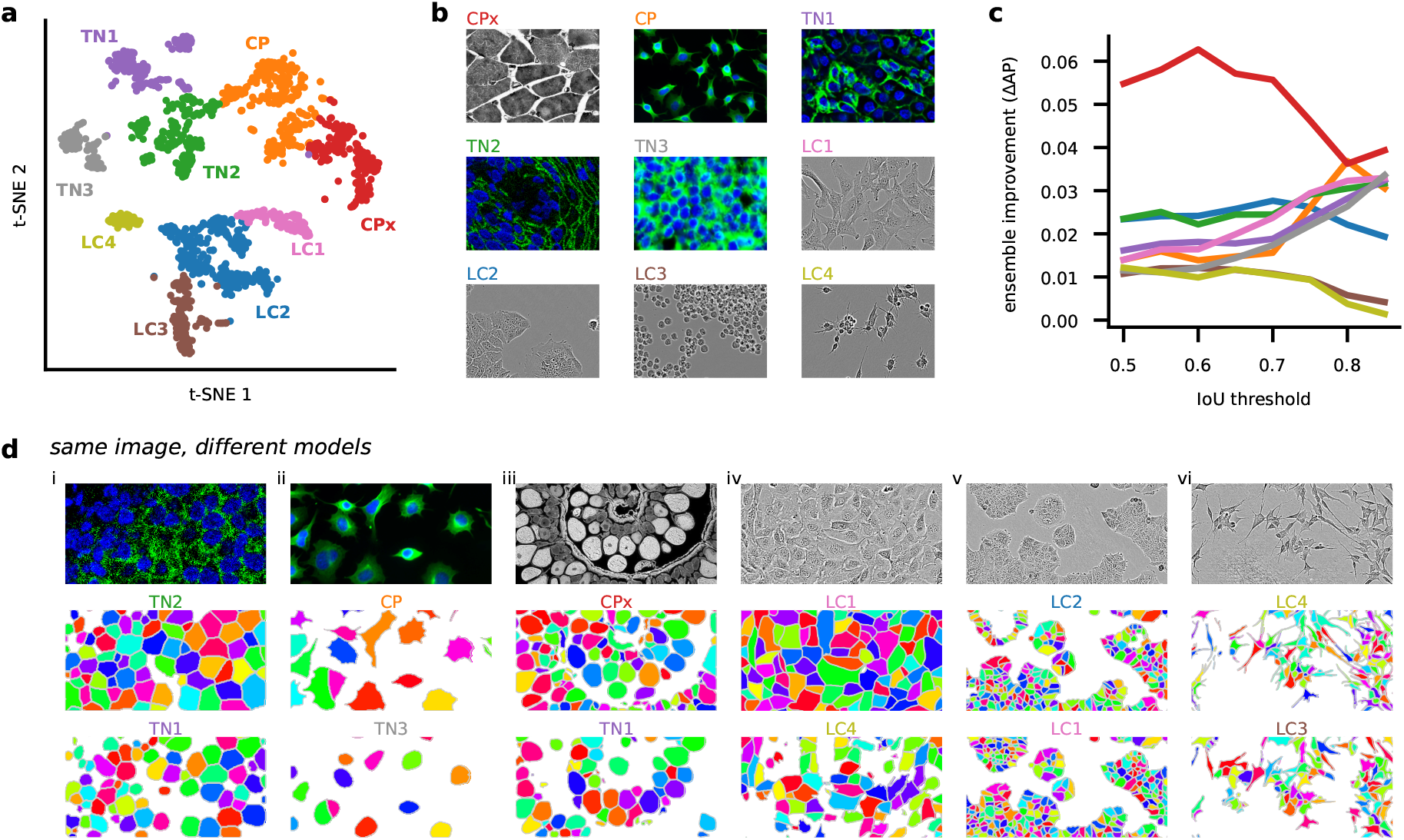
An ensemble of models with different segmentation styles. **a**, t-SNE display of the segmentation styles of images from the Cellpose, LiveCell and TissueNet datasets. The style vector computed by the neural network was embedded in two-dimensions using t-SNE and clustered into 9 groups using the Leiden algorithm. Each color indicated one cluster, with the name chosen based on the most popular image category in the cluster. **b**, Example images from each of the nine clusters corresponding to different segmentation styles. **c**, Improvement of the generalist ensemble model compared to a single generalist model. **d**, Examples of six different images from the test set segmented with two different styles each.

Having obtained nine distinct models, we investigated differences in segmentation style by applying multiple models to the same images (Figure 2d). We saw a variety of effects: the TN1 model drew smaller regions around each nucleus than the TN2 model, which extended the ROIs until they touched each other (Figure 2di); the CP model carefully tracked the precise edges of cells while the TN3 model ignored processes (Figure 2dii); the CPx model segmented everything that looked like an object, while the TN1 model selectively identified only bright objects, assigning dim objects to the background (Figure 2diii); the LC1 model overall identified more cells than the LC4 model, which specifically ignored larger ROIs (Figure 2div); the LC2 model ignored ROIs in very dense regions, unlike the LC1 model which segmented everything (Figure 2dv); the LC4 model tracked and segmented processes over longer distances than the LC3 model (Figure 2dvi); etc. None of these differences are mistakes. Instead, they are different styles of segmenting the same images, each of which may be preferred by a user depending on circumstances. By making these different models available in Cellpose 2.0, we empower users to select the model that works best for them.

For images of nuclei, we did not see an improvement for the ensemble of models compared to a generalist model (Figure S1).

### Cellular segmentation without big data

We have seen so far that segmentation styles can vary significantly between different datasets, and that an ensemble of models with different segmentation styles can in fact outperform a single generalist model. However, some users may prefer segmentation styles not available in our training set. In addition, the ensemble method does not address the out-of-distribution problem, i.e. the lack of generalization to completely new image types. Therefore, we next investigated if a user could train a completely custom model with relatively little annotation effort.

For this analysis, we treated the TissueNet and LiveCell datasets as new image categories, and asked how many images from each category are necessary to achieve high performance. We used as baselines the models shared by the TissueNet and LiveCell teams (“Mesmer” and “LiveCell model”), which were trained on their entire respective datasets. We trained new models based on the Cellpose architecture that were either initialized with random weights (“from scratch”), or initialized with the pretrained Cellpose weights and trained further from there (see also [14]). The diversity of the Cellpose training set allows the pretrained Cellpose model to generalize well to new images, and provides a good starting set of parameters for further fine-tuning on new image categories. The pretraining approach has been successful for various machine vision problems [26–28].

The TissueNet dataset contained 13 image categories with at least 10 training images each, and the LiveCell dataset contained 8. We trained models on image subsets containing different numbers of training images. To better explore model performance with very limited data, we split the 512×512 training images from the TissueNet dataset into quarters. We furthermore trained models on a quarter of a quarter image, and a half of a quarter image. For testing, we used the images originally assigned as test images in each of these datasets.

Figure 3a shows segmentations of four models on the same image from the test set of the “breast vectra” category of TissueNet. The first model was not trained at all, and illustrates the performance of the pretrained Cellpose model. The second model was initialized with the pretrained Cellpose model, and further trained using four 256×256 images from the TissueNet dataset. The third model was trained with 16 images, and the fourth model used all 524 available images. The average precision score for the test image improved dramatically from 0.36 to 0.68 from the first to the second model. Much smaller incremental improvements were achieved for the third and fourth models (0.76 and 0.76). The rapid initial improvement is also seen on average for multiple models trained with different subsets of the data and on all TissueNet categories (Figure 3b). Furthermore, pretrained Cellpose models improved faster than the models trained from scratch: the pretrained model reaches an AP of 0.73 at 426 training ROIs vs 0.68 AP for the model trained from scratch. We also noticed that the pre-trained Cellpose models outperform the strong Mesmer model starting at 1,000 training ROIs, which corresponds to two full training images (512×512). This increase in performance happens despite the Mesmer model being trained with up to 200,000 training ROIs from each image category, and is likely explained by differences between the architecture of the segmentation models.

**Figure 3:**
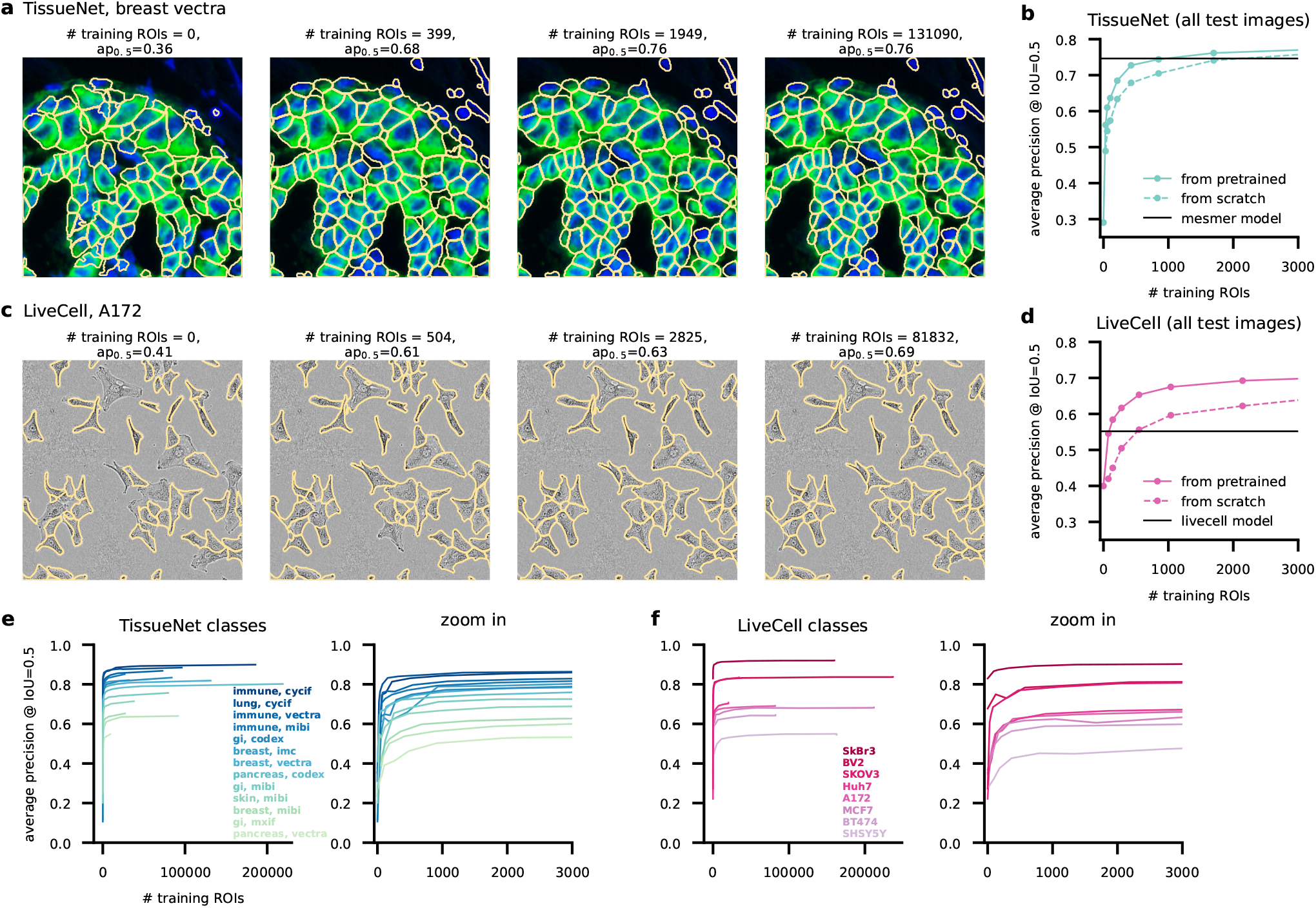
State-of-the-art cellular segmentation does not require big data. **a**, Segmentation of the same test image by models trained with incrementally more images and initialized from the pretrained Cellpose 1.0 model. The image category is “breast, vectra” from the TissueNet dataset. **b**, Average precision of the models as a function of the number of training masks. Shown is the performance of models initialized from the Cellpose parameters or initialized from scratch. We also show the performance of the Mesmer model, which was trained on the entire Tissuenet dataset. **cd**, Same as (a,b) for image category “A172” from the LiveCell dataset. The LiveCell model is shown as a baseline, with the caveat that this model was trained to report overlapping ROIs (see Methods). **e**, (left) Average precision curves for all image categories in the TissueNet dataset. (right) Zoom-in for less than 3,000 training masks. **f**, Same as (e) for the Livecell image categories. **g**, Average of all curves shown in (e,f), as well as the average of the equivalent curves for models initialized from scratch.

We see a similar performance scaling for images from the “A172” category of the LiveCell dataset (Figure 3cd). Performance improves dramatically with 504 training ROIs (equivalent to 2 training images), and then improves much more slowly until it reaches the maximum at 81,832 ROIs. The Cellpose models also appear to outperform the LiveCell model released with the LiveCell dataset [29], although this result should be interpreted cautiously since we evaluated performance after removing mask overlaps which the Cellpose model cannot handle correctly. Finally, we see similar performance scaling across all image categories from both datasets (Figure 3ef). We conclude that 500-1000 training ROIs from each image category are sufficient for near-maximal segmentation accuracy in the TissueNet and LiveCell datasets.

### Fast modelling with a human-in-the-loop approach

We have shown in the previous section that good models can be obtained with relatively few training images. We reasoned that annotation times can be reduced further if we used a “human-in-the-loop” approach [6, 19]. We therefore designed an easy-to-use, interactive platform for image annotation and iterative model retraining. The user begins by running one of the pretrained Cellpose models (e.g. Cellpose 1.0, Figure 4a). Using the GUI, the user can correct the mistakes of the model on a single image and draw any ROIs that were missed or segmented incorrectly. Using this image with ground-truth annotation, a new Cellpose model can be trained and applied to a second image from the user’s dataset. The user then proceeds to correct the segmentations for the new image, and then again retrains the Cellpose model with both annotated images etc. The user stops the iterative process when they are satisfied with the accuracy of the segmentation. In practice, we found that 3-5 images were generally sufficient for good performance. Since only a few images are used for neural network training, the model run times tend to be very short (*<* 1 minute on a GPU).

**Figure 4:**
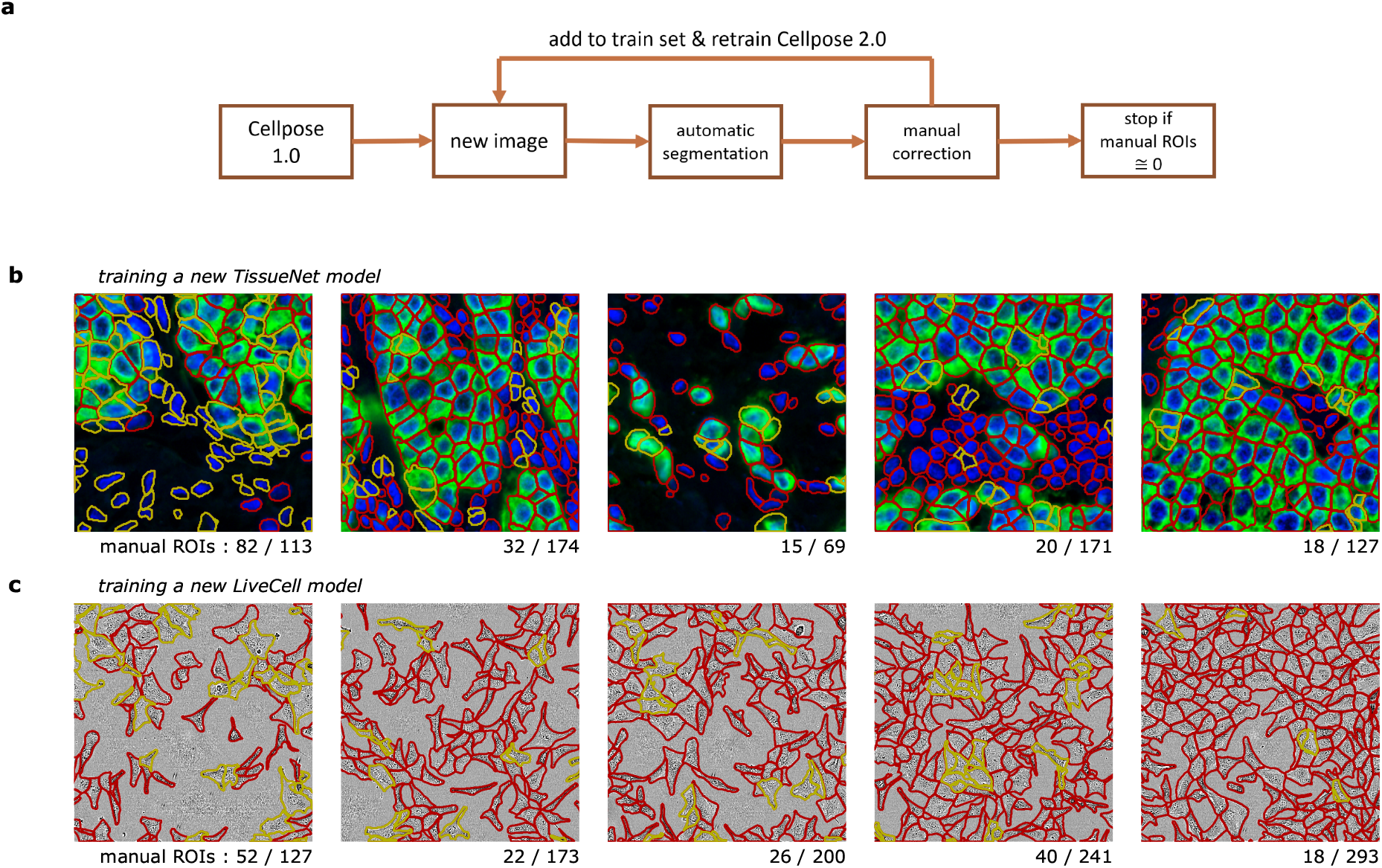
A human-in-the-loop approach for training specialized Cellpose models. **a**, Schematic of human-in-the-loop procedure. This workflow is available in the Cellpose 2.0 GUI. **b**, A new TissueNet model on the “breast, vectra” category was built by sequentially annotating the five training images shown. After each image, the Cellpose model was retrained using all images annotated so far and initialized with the Cellpose parameters. On each new image, the latest model was applied, and the human curator only added the ROIs that were missed or incorrectly segmented by the automated method. The red outlines correspond to cells correctly identified by the model, and the yellow outlines correspond to the new cells added by the human annotator. **c**, Same as (b) for training a LiveCell model on the “A172” category.

To assess the performance of this platform, we trained multiple models with various human-in-the-loop and offline annotation strategies. Critically, we used the same human to train all models, to ensure that the same segmentation style is used for all models. We illustrate two example timelines of the human annotation process (Figure 4bc). For the TissueNet category, the human annotator observed that many cells were correctly segmented by the pretrained Cellpose model, but nuclei without cytoplasm were always ignored, which is likely due to the segmentation style used in the original Cellpose dataset (Figure 1ai). Hence, 82 new ROIs were added and the model was retrained. On the next image, only 32 new ROIs had to be manually added which continued to decrease on the third, fourth and fifth images. Qualitatively, the human annotator observed that the model’s mistakes were becoming more subjective, and were often due to uncertain cues in the image. Nonetheless, the annotator continued to impose their own annotation style, to ensure that the final model captured a unique, consistent style at test time. A similar process was observed for images from the LiveCell dataset (Figure 4c), where 52 out of 127 ROIs had to be drawn manually on the first image, but only 18 out of 293 ROIs had to be drawn on the fifth image.

To evaluate the human-in-the-loop models, we further annotated three test images for each of the two image categories (TissueNet and LiveCell). For comparisons, we also performed complete, offline annotations of the same five training images (from Figure 4bc), and we ran the human-in-the-loop procedure with models either initialized from scratch or from the pretrained Cellpose model. Thus, we could compare four different models corresponding to all possible combinations of online/offline training and pretrained/scratch initialization (Figure 5a). As an upper bound on performance, we annotated the test images twice, with the second annotation performed on images that were mirrored vertically and horizontally (Figure 5b). The average precision between these two annotations can be used as a measure of human accuracy and provides an upper bound for the automated model.

**Figure 5:**
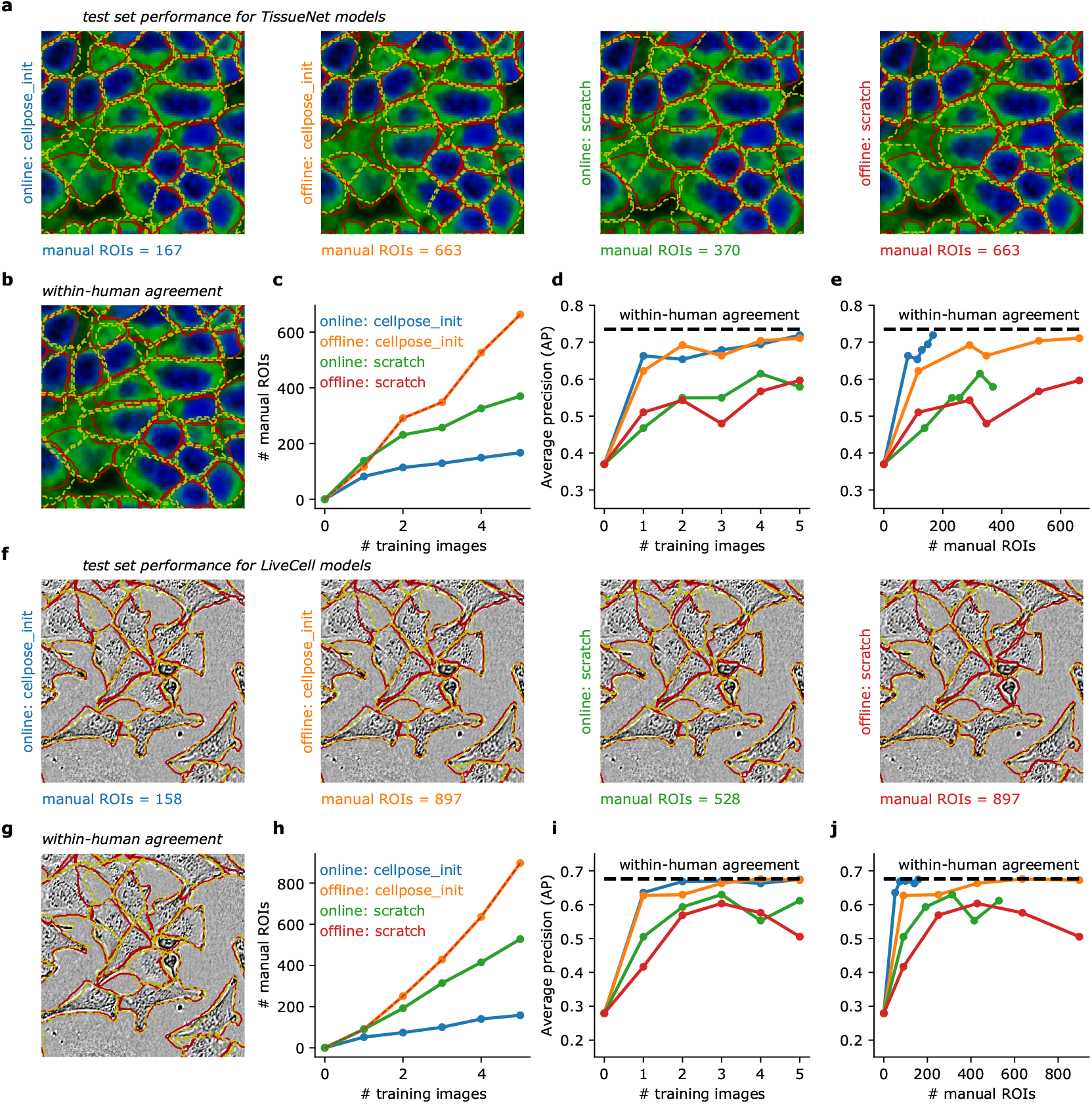
Human-in-the-loop models require minimal human annotation. **a**, Test image segmentations of four models trained on the five TissueNet images from Figure 4b with different annotation strategies. Annotations were either produced with a human-in-the-loop approach (online), or by independently annotating each image without automated help (offline). The models were either pretrained (cellpose init) or initialized from scratch. Red outlines correspond to the ground-truth provided by the same annotator. Yellow outlines correspond to model predictions. **b**, Within-human agreement was measured by having the human annotator segment the same test images twice. For the second annotation, the images were mirrored horizontally and vertically to reduce memory effects. **c**, Total number of manually-segmented ROIs for each annotation strategy. **d**, Average precision at an IoU of 0.5 as a function of the number of training images. **e**, Average precision curves as a function of the number of manually annotated ROIs. **f-j**, Same as (a-e) for the image category “A172” from the LiveCell dataset. All models were trained on the images from Figure 4b, with the same annotation strategies.

The online models in general required fewer manual segmentations than the offline models (Figure 5c). Furthermore, the online model initialized from Cellpose required many fewer manual ROIs than the online model initialized from scratch. Overall, we only needed to annotate 167 total ROIs for the online/pretrained model, compared to 663 ROIs for a standard offline approach. Performance-wise, models pretrained with the standard Cellpose dataset did much better than models initialized from scratch (Figure 5c). Of the four models, the online/pretrained model was unique in achieving near-maximal precision with very few manual ROIs (Figure 5e). All of these results were confirmed with a different set of experiments on a LiveCell image category (Figure 5f-j). In both cases, 100-200 manually-segmented ROIs were sufficient to achieve near-maximal accuracy, and the process only required 1-2 hours of the user’s time.

## Discussion

Here we have shown that state-of-the-art biological segmentation can be achieved with relatively little training data. To show this, we used two existing large-scale datasets of fluorescence tissue images and phase-contrast images, as well as a new human-in-the-loop approach we developed. We are releasing the software tools necessary to run this human-in-the-loop approach as a part of the Cellpose 2.0 package. Finally, we showed that multiple large datasets can be used to generate a zoo of models with different segmentations strategies, which are also immediately available for Cellpose users.

Our conclusions may seem at odds with the conclusions of the original papers introducing the large-scale annotated datasets. The TissueNet authors concluded that performance saturated at 10^4^–10^5^ training ROIs. The LiveCell authors concluded that segmentation performance continued to increase when adding more training data. The discrepancy with our results may be due to several factors. First, we found that models initialized with Cellpose saturated their performance much more quickly than models trained from scratch. Second, Cellpose as a segmentation model appeared to perform better than both the Mesmer and LiveCell models, and this in turn may lead to higher efficiency in terms of required training data. Third, we focused on the initial portion of the performance curves where models were trained on only tens to hundreds of ROIs, which was below the first few datapoints considered in the TissueNet and LiveCell studies, even splitting images into quarters to explore very limited training data scenarios. Fourth and finally, we used a large set of image augmentations to further increase the diversity of the training set images and improve generalizability [5].

Our analysis also showed that there can be large differences in segmentation style between different annotators, even when their instructions are the same. This variability hints at a fundamental aspect of biological segmentation: there are often multiple “correct” solutions, and a biologist may prefer one segmentation style over another depending on the purpose of their study. The goal of having a single, global generalist algorithm for biological segmentation is therefore unachievable.

Future efforts to release large annotated datasets should focus on assembling highly-varied images, potentially using algorithms to identify out-of-distribution cell types [30, 31], and should limit the number of training exemplars per image category. We renew our calls for the community to contribute more varied training data, which is now easy to generate with the human-in-the-loop approach from Cellpose 2.0.

## Acknowledgments

This research was funded by the Howard Hughes Medical Institute at the Janelia Research Campus. We thank the authors of [6] and [7] for making their datasets and code publicly available.

## Data and code availability

The code and GUI are available at https://www.github.com/mouseland/cellpose.

## Methods

The Cellpose code library is implemented in Python 3 [32], using pytorch, numpy, scipy, numba, opencv [20, 33–36]. The graphical user interface additionally uses PyQt, pyqtgraph and scikit-image [37–39]. The figures were made using matplotlib and jupyter-notebook [40, 41].

### Models and training

#### Cellpose model

The Cellpose model is described in detail in [5]. Briefly, Cellpose is a deep neural network with a U-net style architecture and residual blocks [42, 43]. Cellpose predicts 3 outputs: the probability of a pixel being inside a cell (1), the flows of pixels towards the center of a cell in X (2) and Y (3). The flows are then used to construct the cell ROIs. The Cellpose default model (“cyto”) was trained on 540 images of cells and objects with 1 or 2 channels (if the image had a nuclear channel). This is the pretrained model used, which we refer to as the “Cellpose 1.0” model.

#### Training

All training was performed with stochastic gradient descent. In offline mode, the models, either from pretrained or from scratch, were trained for 300 epochs with a batch size of 8, a weight decay of 0.0001 and a learning rate of 0.1. The learning rate increased linearly from 0 to 0.1 over the first 10 epochs, then decreased by factors of 2 every 5 epochs after the 250th epoch. There were a minimum of 8 images per epoch, so if fewer than 8 images were in the training set then they were randomly sampled with replacement to create a batch of 8 images. In online mode, training occurred for only 100 epochs, otherwise the parameters were the same. The learning rate was again increased linearly from 0 to 0.1 over the first 10 epochs, but no annealing of the learning rate occured toward the end of training. We observed slight performance improvements for the models trained from scratch but not from pretrained for 300 epochs of training compared to 100 epochs.

In Figure 3, we trained on subsets of images in the training set, from 0.25 (a quarter image), 0.5 (a half image), 1, 2, 4, in powers of 2 up to 2048 depending on the number of images in the cell class. We trained at each of these subset sizes 5 times with 5 different random subsets of images and averaged the performance and the number of ROIs used for training across these 5 networks.

The generalist and ensemble models in Figure 2 and Figure S1 were trained from scratch for 500 epochs with a batch size of 8, a weight decay of 0.00001 and a learning rate of 0.2. The learning rate increased linearly from 0 to 0.2 over the first 10 epochs, then decreased by factors of 2 every 10 epochs after the 400th epoch. The model used to compute style vectors in Figure 2a was trained with images sampled from the Cellpose “cyto” dataset, the TissueNet dataset, and the LiveCell dataset, with probabilities 60%, 20% and 20% respectively. The generalist model that was compared to the ensembles (Figure 2c) was trained with images sampled from the style vector clusters with equal probabilities. The ensemble models were trained using all the training images classified in the cluster with equal probability.

For all training, images with fewer than 5 ROIs were excluded.

#### Style clustering and classification

In Cellpose, we perform global average pooling on the smallest convolutional maps to obtain a representation of the “style” of the image, a 256-dimensional vector [12, 23, 44]. For the clustering of style vectors in Figure 2a we used all of the Cellpose “cyto” training images, 20% of the TissueNet training images, and 20% of the LiveCell training images. We then ran the Leiden algorithm on these style vectors with 100 neighbors and resolution 0.45 to create 9 clusters of images [24]. For the images in the training set not used for clustering and in the test set, we used a K-nearest neighbor classifier with 5 neighbors to get their cluster labels.

For the clustering in Figure S1a we used all of the training images in the Cellpose “nuclei” dataset. We then ran the Leiden algorithm on these style vectors with 50 neighbors and resolution 0.25 to create 6 clusters of images. For the images in the test set, we used a K-nearest neighbor classifier with 5 neighbors to get their cluster labels.

#### Evaluation

For all evaluations, the flow error threshold (quality control step) was set to 0.4. When evaluating models on test images from the same image class (Figure 3), the diameter was set to the average diameter across images in the training set. For the online/offline comparisons in Figure 4 and Figure 5 the diameter was set to 18 for all the breast vectra TissueNet images and 34 for all the A172 LiveCell images, which was their approximate average diameter in the training set. When evaluating the ensemble versus generalist model performance (Figure 2 and Figure S1), the diameter was set to the diameter of the given test image for all models, so that we can rule out error variability due to imperfect estimation of object sizes.

### Model comparisons

We compared the performance of the Cellpose models to the Mesmer model trained on TissueNet [6] and the anchor-free model trained on LiveCell [7, 29].

#### Mesmer model

We used the Mesmer-Application.ipynb notebook provided in the DeepCell-tf github repository to run the model on the provided test images with image mpp=0.5 and compartment=“whole-cell” [6, 45].

#### Livecell model

We used the pretrained Livecell anchor-free model provided by the authors to run the model on the provided test images [29, 46]. The ROIs returned by the algorithm could have overlaps, and therefore we removed the overlaps as described in the Livecell Dataset section below.

#### Quantification of segmentation quality

We quantified the predictions of the algorithms by matching each predicted mask to the ground-truth mask that is most similar, as defined by the intersection over union metric (IoU). Then we evaluated the predictions at various levels of IoU; at a lower IoU, fewer pixels in a predicted mask have to match a corresponding ground-truth mask for a match to be considered valid. The valid matches define the true positives *TP*, the ROIs with no valid matches are false positives *FP*, and the ground-truth ROIs which have no valid match are false negatives *FN*. Using these values, we computed the standard average precision metric (*AP*) for each image:

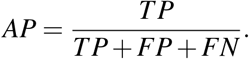

The average precision reported is averaged over the *AP* for each image in the test set.

### Datasets

#### TissueNet

The TissueNet dataset consists of 2601 training and 1249 test images of 6 different tissue types collected using fluorescent microscopy on 6 different platforms, and each image has manual segmentations of the cells and the nuclei (https://datasets.deepcell.org/) [6]. We only used the cellular segmentations in this study. We excluded the “lung mibi” type from Figure 3 because it only contained 1 training image and 4 test images. We thus used the other 13 types: pancreas codex, immune cycif, gi mibi, lung cycif, gi codex, breast vectra, gi mxif, skin mibi, breast mibi, immune vectra, breast imc, immune mibi, and pancreas vectra. The training images are 512 × 512 pixels. To enable subsets consisting of fewer ROIs in Figure 3, we divided each training image into 4 parts and used those in the training protocol.

#### LiveCell

The LiveCell dataset consists of 3188 training and 1516 test images of 8 different cell lines collected using phase-contrast microscopy, and each image has manual segmentations of the cells (https://sartorius-research.github.io/LIVECell/) [7]. The 8 cell lines were MCF7, SkBr3, SHSY5Y, BT474, A172, BV2, Huh7, and SKOV3. The images were segmented with overlaps allowed across ROIs. The Cellpose model cannot predict overlapping ROIs, therefore the overlapping pixels were reassigned to the mask with the closest centroid. ROIs with more than 75% of their pixels overlapping with another ROI were removed. These non-overlapping ROIs were used to train Cellpose and benchmark the results.

#### Cellpose cyto dataset

This dataset was described in detail in [5]. Briefly, this dataset consisted of 100 fluorescent images of cultured neurons with cytoplasmic and nuclear stains obtained from the CellImageLibrary [47]; 216 images with fluorescent cytoplasmic markers from BBBC020 [48], BBBC007v1 [49], mouse cortical and hippocampal cells expressing GCaMP6 using a two-photon microscope, and 10 images from confocal imaging of mouse cortical neurons with cytoplasmic and nuclear markers, and Google image searches; 50 images taken with standard brightfield microscopy from OMERO [50], and google image searches; 58 images where the cell membrane was fluorescently labeled from [51] and google image searches; 86 images from microscopy samples which were either not cells, or cells with atypical appearance from google image search; 98 non-microscopy images of repeating objects from google image search.

#### Cellpose nucleus dataset

This dataset was described in detail in [5]. Briefly this dataset consisted of images from BBBC038v1 [52, 53], BBBC039v1 [52], MoNuSeg [54], and ISBI 2009 [55].

---

**S1:**
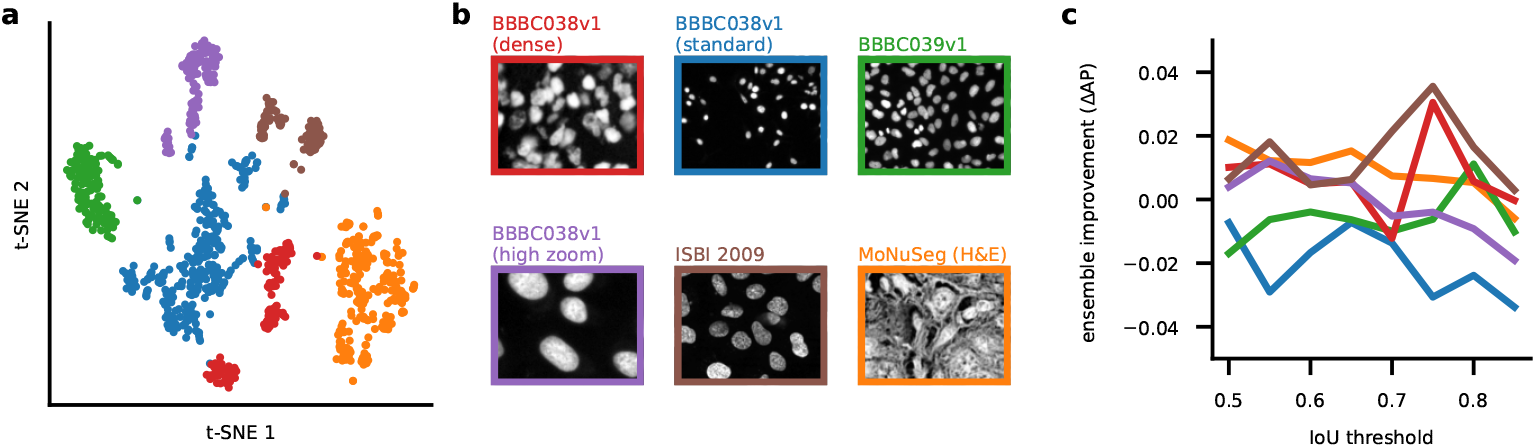
Specialization of the pretrained model for images of nuclei. **a**, t-SNE embedding of segmentation styles for each image, colored acorrding to cluster identity. **b**, Representative example images from each class. **c**, AP improvement of the model ensemble over a single generalist model.

